# Foraging guild structure of seabirds

**DOI:** 10.1101/2023.12.09.570957

**Authors:** Juan Hernández, Jose Ignacio Arroyo

## Abstract

An ecological guild is a group of species that exploit the same resources, or that exploit the same or different resources in a related way. We built The Foraging Guilds of Seabirds database (FGSdb) by compiling a global database of 311 seabird species (from a total of 346 known) and assigning to each of them their diet types and foraging strategy. Across all seabirds, there were found 22 diets and 30 strategies. The number of diet categories for a species varied between 1 and 11, and the number of strategies varied from 1 to 9, with averages of 2.71 and 3.65, being the ratio diet/strategies of 0.74 (∼3/4), meaning that on average with four strategies they can exploit up to 3 diet items. Beyond this description, we show that the Gusein-Zade model fits well both the frequency rank and number of species per guild distributions. Our database and analyses provide a useful resource database for future studies and demonstrate simple rules behind guild structure.

## Introduction

Understanding ecological functional groups is relevant given that their structure ultimately affects the functioning and stability of ecosystems. Within functional groups, particularly ecological guilds, are groups of species that exploit the same resources, or that exploit different resources in a related way. They do not necessarily belong to the same niche or are phylogenetically related, but for practical reasons often are. However, they do match the general concept of functional groups as exploiting the same resources can perform the same or highly similar ecosystem function (Simberloff & Dayan 1991, Wilson 1999).

Often, ecological guilds are studied by identifying the dietary categories that species use or the different behaviors or strategies they use to get those diets and analyzing the composition and structure (Uetz et al. 1999, Cardoso et al. 2011, Kissling et al. 2014, Collard et al. 2021). Composition refers to for which and how many species a given guild is composed, and structure commonly refers to how is composed and whether each of the groups differs in their abundance or whether species conform to different groups based on their properties. This is often analyzed by classifying species through a multidimensional scaling or clustering method (Uetz et al. 1999, Cardoso et al. 2011, Collard et al. 2021).

Guilds of birds have fundamental ecological importance as they participate in ecosystem services (that include seed dispersal, and pollination), environmental management, and conservation. Despite the guild structure of terrestrial bird communities has been characterized (De Graff et al. 1985, González-Salazar et al. 2014), the guild of seabirds at a global scale remains unstudied but restricted to particular communities or specific taxa. Seabirds are a functional group and refer to all birds that live in the ocean environment, including marine and coastal. Seabirds are a paraphyletic group (including the orders Procellariiformes, Charadriiformes, Sphenisciformes, Pelecaniformes, Phaethontiformes, and Suliformes) and have independently and convergently acquired different adaptations to pelagic and aquatic life (Hackett et al. 2008). The study of their ecology (Oro et al. 2004) particularly diet and foraging (Shealer 2002, de L. Brooke 2004) is a relevant topic of study due to their relationship with humans, role in fisheries (Montevecchi 2002), and indicators in marine environments (Parsons et al. 2008).

Here we characterize the unstudied structure of the foraging guilds of seabirds. We describe both the strategies and diets species use and provide a hierarchical classification that could be a useful resource for future studies. Beyond this description, we show that the probability distributions that describe their structure are explained by a simple model, the Gusein-Zade function.

## Methods

### Data

We built The Foraging Guilds of Seabirds database (FGSdb; Hernandez and Arroyo 2023; available at https://doi.org/10.5281/zenodo.10258241) by compiling data on dietary niche breadth and prey capture flexibility were manually obtained from the Handbook of the Birds of the World (https://birdsoftheworld.org/bow/home). We used as an initial reference for diet guilds the classification of foraging guilds of North American breeding birds (De Graaf et al. 1985; González-Salazar et al.2014). Secondly, for those species for which we did not find data or a single strategy or diet item, we searched for data directly from the literature. Statistical analysis was performed in the R environment (R Core Team 2021).

### Analysis

We analyzed the frequency rank distribution (Brookes and Griffiths 1978, Newman 2005) and the number of species per guild distribution (analogous to the number of species in a genus; Willis 1922, Yule 1924, Simon 1955). Briefly, going from a probability density function to a frequency rank distribution can be done by equating the cumulative distribution function (cdf) to a reversed normalized rank (Fontanelli et al. 2016). We estimated bin size as *b* = *n*^½^. To assess any effect of the bin size on the form of the distributions we also used Scott’s rule, as W=3.49sn^-⅓^, where W is bin size, s is the standard deviation of the data, and n is the number of observations in the sample, and the number of bins as b=max-min/W, where b is the number of bins, *max* is the maximum value, *min* the minimum value. We fitted frequency ranks and species per guild distributions to the Gusein-Zade model using a simple linear regression as implemented in R (R Core Team 2021).

## Results

### Database description

Briefly, we built a relational database with a first file containing the species and their strategies and diets affiliations, as a vector. In a second file, we put the strategies and their classification, and in a third file, we put the diets and their classification (Fig. S1). For a total of 311 seabird species (from a total of 346 known, BirdLife International 2012), we defined a total of 30 strategies and 22 diets. Each of these groups was subcategorized into larger groups.

The definition of the relevant foraging strategies is as follows (Ainley 1977, Duffy 1980, Duffy 1983a, Duffy 1983b, Duffy & Jackson 1986, Burger 1988, Marine 1988, Croxall & Cooper 1985, Camphunysen and Garthe 2004): *Scooping*: Swimming birds, scooping up small prey from just below the surface. *Wading, filtering foraging:* wading, filtering, or gathering, in shallow areas. *Ground forager:* a species that forage on the ground. *Probing:* collector in deep areas. *Surface seizing:* birds settle on the surface and grasp food with the bill. *Surface picking:* birds swim pecking small prey at the surface. *Deep plunging:* aerial seabirds diving into the sea and completely disappearing underwater. *Shallow plunging:* aerial seabirds diving into the sea and partly disappearing underwater. *Pursuit plunging:* aerial seabirds plunging into the water and continuing with an underwater pursuit. *Pursuit diving:* swimming seabirds that perform deep dives and are known to search for prey in an underwater pursuit or search for prey at the bottom. *Scavenging:* search for discarded items or food. *Pattering:* fly low over the surface of the water, zig-zag movement, and strike the surface of the water with your feet while still in the air. *Hydroplaning:* low flight over the surface, filtering surface layers. *Kleptoparasitism:* steal of prey. *Contact dipping*: bird in flight pecks food from or just below the surface. Only bill, head, or breast make contact with water. *Hoever dipping:* seabird floats in the air in contact with the water capturing prey. *Dipping:* aerial seabirds making repeated dives while hardly touching the water (remaining airborne) and picking up small prey. *Aerial pursuit:* individuals pursue prey in the air. *Shallow dives:* bird submerges completely but penetrates < 0.5 m and swims very little to catch food. *Wing:* captures prey through wing movement. *Surface plung:* shallow booby. *Kick splashing:* individuals kick trough the surface while splashing. *Battering or drowning:* other seabirds individuals due to drowning or blows. *Plunge diving:* diving without pursuit. *Skimming:* low flight over the water surface, touching the surface with the beak. *Walking:* birds capture prey by walking on the surface. *Ships follow:* persecution of ships. *Cannibalism:* eat individuals of the same species.

These strategies were grouped into five groups depending on the space niche they use, marine, superficial, or aerial (Ashmole 1971, Table S1).

The definition of diets is straightforward. The higher-level definitions need some description. Each of these diet types was grouped into trophic groups; parasites, herbivores, primary, secondary, and tertiary/quaternary consumers. For example, species feeding on algae and terrestrial plants collapsed into herbivores (Table S1).

### Statistics

The number of strategies they had ranged from 1 to 9, with an average of 2.71. The number of diet categories to which a species belongs ranges from 1 to 11, with an average of 3.75. The ratio diet/strategies of 0.74 (∼3/4), meaning that on average with four strategies they can exploit up to three diet items. The most frequent among seabird species were scavenging pursuit diving (or bottom feeding) and surface seizing and the less diverse were battering or drowning ship follows and cannibalism (Fig. 1A). The most frequent diets were fish, crustaceans, cephalopods, and the least diverse were chaetognaths, lampreys, and polychaetes (Fig. 1B).

**Figure 1.**
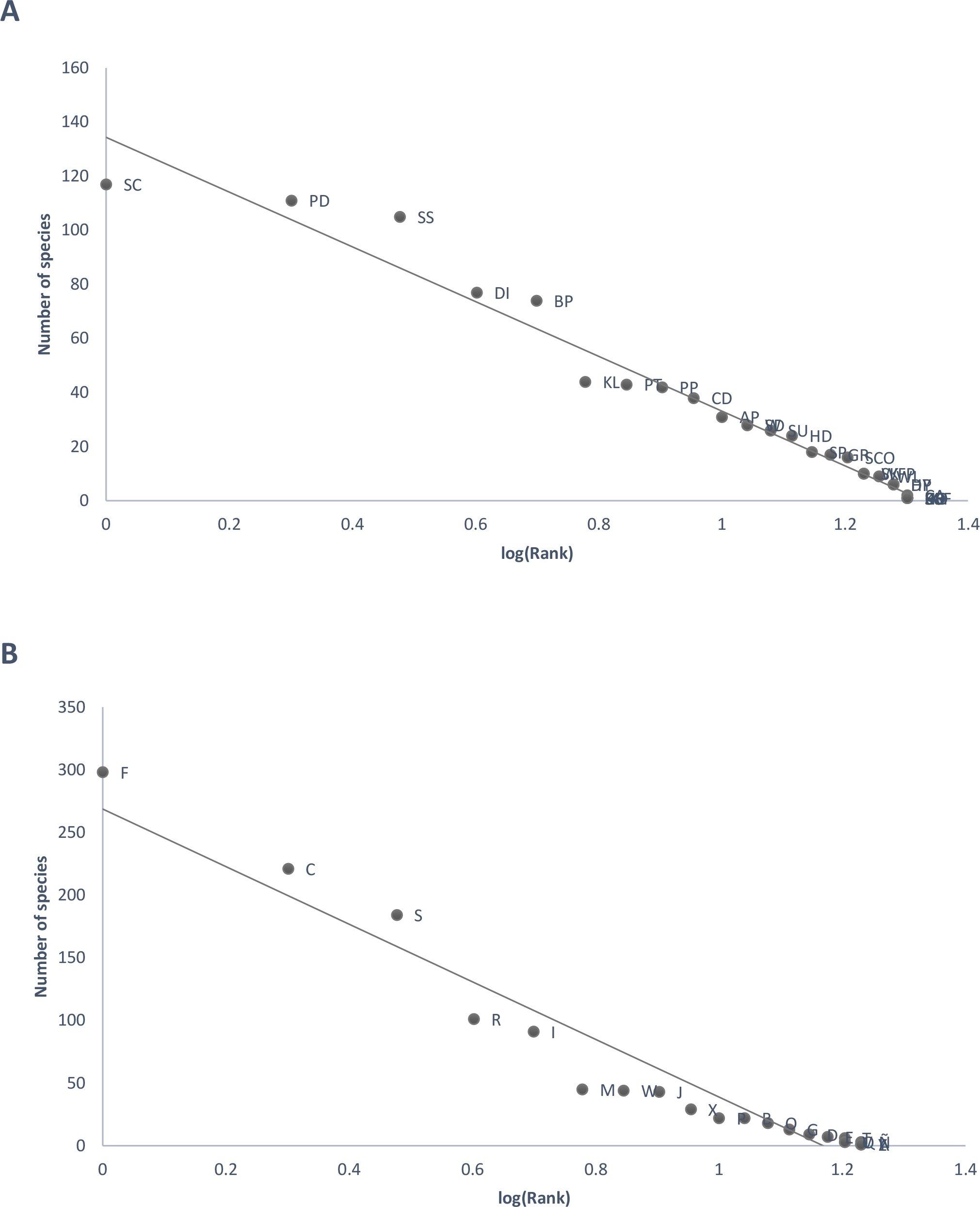
Frequency-rank relationships of guilds of strategies and diet types. A) strategies, B) diet types. AP: Aerial pursuit, HD: Hoever dipping, HY: Hydroplaning, PT: Pattering, SU: Surface plung, W: Wing, DI: Dipping, DP: Deep plunging, PD: Pursuit diving, PP: Pursuit plunging, SP: Shallow plunging, CA: Aerial capture, BP: Plunge diving, PR: Probing, SD: Shallow dives, CB: Cannibalism, KL: Kleptoparasitism, SCF: Ships follow, BD: Battering or drowning, CD: Contact dipping, GR: Ground forager, KS: Kick splashing, SC: Scavenging, SCO: Scooping, SG: Surface picking, SK: Skimming, SS: Surface seizing, WFP: Wading, filtering foraging, WL: Walking, HO: Hovering; F: Fish, O: Reptiles, S: Cephalopods, X: Mammals, B: Amphibia, J: Seabirds, Y: Sponge, M: Mollusk, A: Polychaete, C: Crustacean, E: Echinoderm, W: Worm, Ñ: Chaetognaths, D: Hydrozoa, T: Tunicates, I: Insects, G: Plankton, V: Algae, P: Terrestrial plant, Z: Cannibal, R: Scavenger, Q: Opportunistic feeder, L: Lamprey. Both relationships fit significantly into the Gusein-Zade mode as in Eq. 1.

### Model fitting

Beyond this description, we provide statistical modeling for the frequency rank and number of species per guild distributions. After examination of the data, we noticed that these three relationships fit well with the Gusein-Zade function (Gusein-Zade 1988, see also Li et al. 2010 for an explanation of the origin of the distribution),

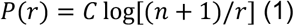

Where P is the frequency, r is rank, and n (which denotes the number of items to be ranked) and C are constants. Eq. 1 can be expressed in log scales as,

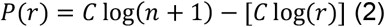

Note that Eq. 2 can take the form of a linear model; P(r)=a-Cx, where a=C log(n+1), x=log(r), then if we plot P(r)-log(r) there should be a linear pattern. Importantly, these plots are easy to interpret given that the y-axis is in arithmetic scale and the x-axis is the rank, but in log scale.

Both the rank distributions (Fig.1) and number of species in guild distribution (Fig. 2) had a significant fit to Eq. 2 (Table 1). We fitted data separately for the distribution of species in guilds of strategies and diets. In these two cases, the slope was ∼-7, but when doing the fit for all together the slope was the double, ∼-16. The intercept had a similar trend.

**Table 1.**
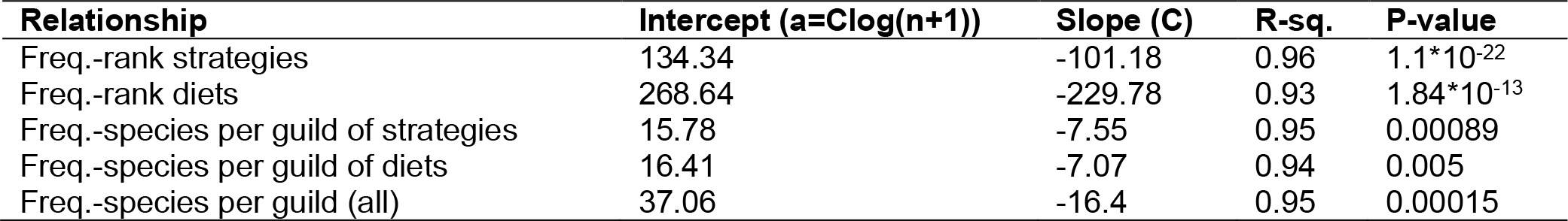
Estimated parameters for the Gusein-Zade model.

| Relationship | Intercept ($a=C\log(n+1)$) | Slope (C) | R-sq. | P-value |
| --- | --- | --- | --- | --- |
| Freq.-rank strategies | 134.34 | -101.18 | 0.96 | $1.1 \cdot 10^{-22}$ |
| Freq.-rank diets | 268.64 | -229.78 | 0.93 | $1.84 \cdot 10^{-13}$ |
| Freq.-species per guild of strategies | 15.78 | -7.55 | 0.95 | 0.00089 |
| Freq.-species per guild of diets | 16.41 | -7.07 | 0.94 | 0.005 |
| Freq.-species per guild (all) | 37.06 | -16.4 | 0.95 | 0.00015 |

**Figure 2.**
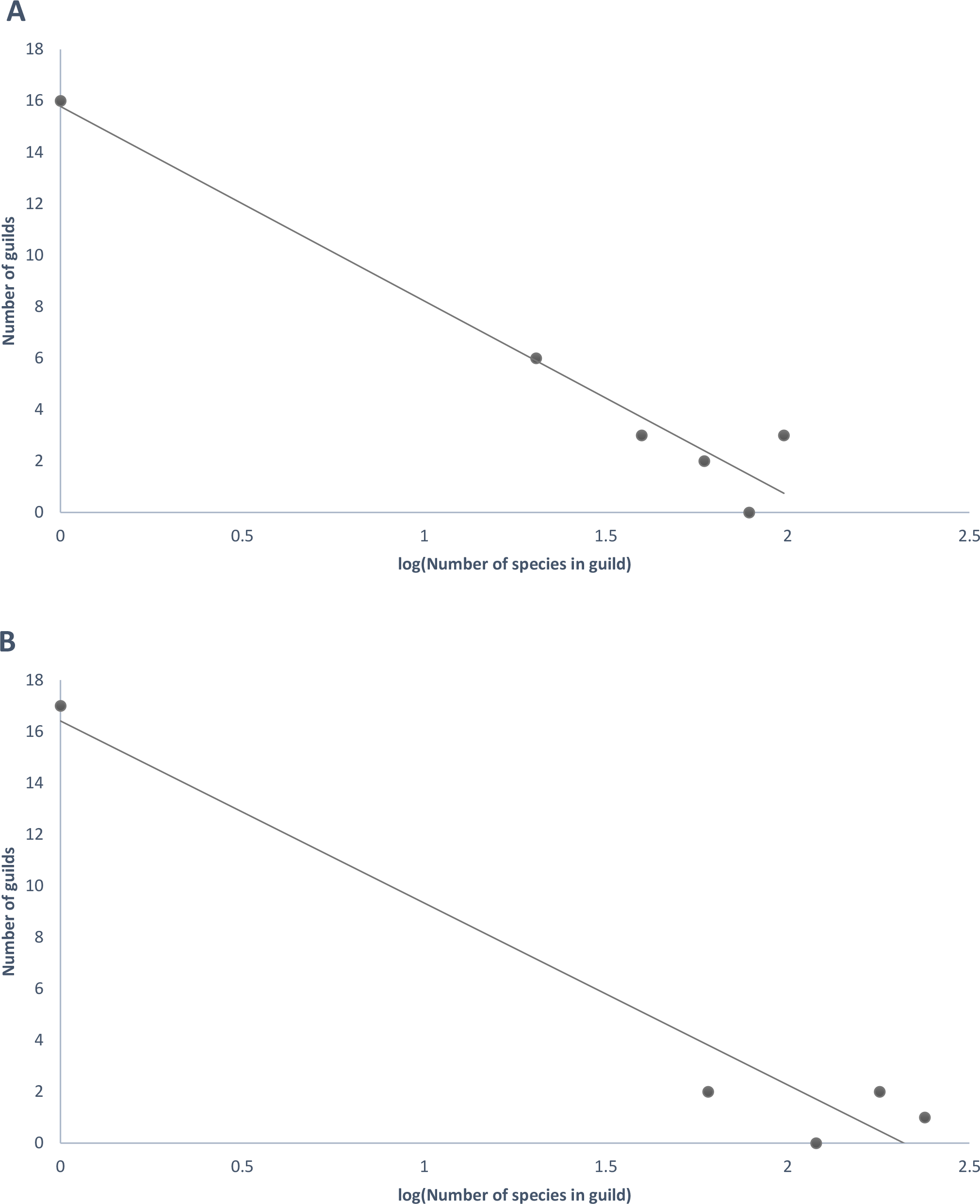
Distribution of number of species per guild. The distribution fits significantly to the Gusein-Zade model as in Eq. 1. Values in the y-axis correspond to lower limits of bin intervals.

## Discussion

This database has different utilities, such as being a reference for studies at a local scale, and that seek for example, to understand the redundancy (and subsequently other properties such as ecosystem functioning) of a community, or to use the communities as ecological indicators of other features of the ecosystem (O’Connell et al. 2000).

We based our definition of guilds mostly on previous classifications of terrestrial birds (De Graf et al. 1985, González-Salazar et al. 2014) and there are common guilds in both the terrestrial and marine environment, such as species that eat fish, arthropods, etc., but there are differences. For example, in terrestrial environments granivores are common, but absent in marine, and on the other hand cephalopods are absent in terrestrial environments. In comparison with previous studies, here we took a step forward and defined a higher level of guild structure for both strategies and diets.

An interesting pattern is the average number of behaviors used per diet or the relationship between behaviors and diet. In this system, this could indicate a benefit/cost ratio, i.e., how many strategies do species need to evolve to be able to exploit a given number of resource items? In this case, the ratio diet/strategies of 0.74 (∼3/4), meaning that on average with three strategies they can exploit up to 4 diet items.

The number of elements in a group and their counterpart, the frequency rank, are a pair of highly studied patterns in biology. For example, classic studies include the distributions of the number of species in a genus (Willis and Yule 1922, Yule 1924) that have in most cases a very well-defined power-law distribution. Often, the structure of ecological groups, or particularly the structure of guilds has been studied using descriptive approaches. Here we found that the frequency-rank distribution and number of species per guild follow the same distribution, the Gusein-Zade function. This is a good example of how complex systems such as letters in a language or guilds in a taxonomic group respond to the same model. Despite differences in the particularities of the system, there could be generic processes of birth and death of new letters or in this case, new guilds that give origin to macroscopic patterns that have a predictable pattern (Elliott-Graves 2022).

Beyond this description and patterns that we found, some alternative approaches that could be further explored include analyzing the relationship between the number of species and guilds, and the relationship between the number of diets and strategies. Also, it would be interesting to make a bipartite network of species and strategies/diets, or a network based on a matrix of presence/absence of strategies and diets across all species. Moreover, a critical question is whether this pattern is the same not just for seabirds but for all birds or wider taxonomic groups, and also for other guilds or ecological groups.

## Conclusion

Understanding the functional structure of communities is important given its impact on ecosystem functioning and stability. Here we developed The Foraging Guild of Seabird database (FGSdb), including data on both the strategies and diets that seabird species have, i.e. how and what resources species use. The most remarkable finding was that the rank and number of species per guild fitted both well to the Gusein-Zade model, a logarithmic function that was originally formulated for the distribution of letters in the Russian language. Beyond the development of this database, which could have several applications, our study shows that the structure of ecological guilds has a predictable structure. This study opens the question of whether other ecological groups and taxa have a predictable group structure which are the models that better explain their structure and whether there is a universal model for all groups and taxa.

## Acknowledgments

JIA acknowledges a Beca de Doctorado Nacional (21130515) from the Agencia Nacional de Investigacion y Desarrollo (ANID) and USA NSF projects: ‘Building and Modeling Synthetic Bacterial Cells’ (1840301), and ‘Towards a unified theory of regulatory functions and networks across biological and social systems’ (2133863). Center for Mathematical Modeling (CMM), FB210005, BASAL funds for centers of excellence from ANID-Chile.

