## Supplementary Material for "Foraging guild structure of seabirds"

### Supplementary Material for
### Foraging guilds structure of seabirds

Juan Hernández* (1, a), Jose Ignacio Arroyo* (2, 3, b)

(1) Department of Ecology, Pontifical Catholic University of Chile, Santiago Chile
(2) Santa Fe Institute, New Mexico, USA
(3) Center for Mathematical Modeling, Santiago, Chile

*All authors contributed equally
(a)
(b)

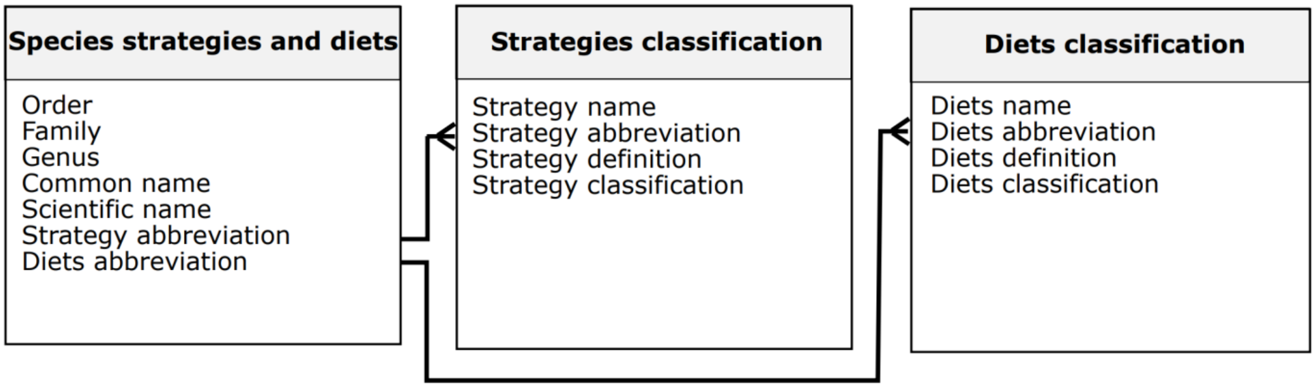


**Figure 1. Schema of the FGSdb database.** Each of the boxes represents a tsv file, and each of the listed titles is a column in that file.

Table 1. Classification of strategies and diets guilds.

| **Diet classification** | **Abb** | **Diets** | **Abb.** |
|---|---|---|---|
| Tertiary/Quaternary consumer | TERQUA | Fish | F |
| | | Reptiles | O |
| | | Cephalopods | S |
| | | Mammals | X |
| | | Amphibia | B |
| | | Seabirds | J |
| | | Sponge | Y |
| | | Mollusk | M |
| | | Polychaete | A |
| | | Crustacean | C |
| | | Echinoderm | E |
| | | Worm | W |
| | | Chaetognaths | Ñ |
| | | Hydrozoa | D |
| | | Tunicates | T |
| Secondary consumer | SEC | Insects | I |
| | | Plankton | G |
| Primary consumer/ Herbivore | HER | Algae | V |
| | | Terrestrial plant | P |
| Opportunistic | OPD | Cannibal | Z |
| | | Scavenger | R |
| | | Opportunistic feeder | Q |
| Parasitic | PAR | Lamprey | L |
| **Strategy classification** | | **Strategies** | |
| Aerial | AE | Aerial Pursuit | AP |
| | | Hoever Dipping | HD |
| | | Hydroplaning | HY |
| | | Pattering | PT |
| | | Surface Plung | SU |
| | | Wing | W |
| Aerial/Diving | AERDIV | Dipping | DI |
| | | Deep Plunging | DP |
| | | Pursuit Diving | PD |
| | | Pursuit Plunging | PP |
| | | Shallow Plunging | SP |
| | | Aerial capture | CA |
| Diving | DIV | Plunge Diving | BP |
| | | Probing | PR |
| | | Shallow Dives | SD |
| Opportunistic | OPS | Cannibalism | CB |
| | | Kleptoparasitism | KL |
| | | Ships Follow | SCF |
| Superficial | SUP | Battering or drowning | BD |
| | | Contact Dipping | CD |
| | | Ground Forager | GR |
| | | Kick Splashing | KS |
| | | Scavenging | SC |

| | | | |
|---|---|---|---|
| | | Scooping | SCO |
| | | Surface Picking | SG |
| | | Skimming | SK |
| | | Surface seizing | SS |
| | | Wading,<br>filtering foraging | WFP |
| | | Walking | WL |
| | | Hovering | HO |

**Text 1.**

**What do seabirds' diet categories feed on.**
Fish: omnivorous
Lamprey: parasites (blood feeders), predators (flesh feeders), parasite-predators (blood-and-flesh feeders), and scavengers or carrion feeders
Reptiles: omnivorous
Cephalopods: cephalopods are mollusks. They are predators
Mammals: omnivorous
Amphibia: omnivorous
Seabirds: omnivorous
Crustacean: omnivorous
Insects: most are herbivorous, feeding on seaweed
Echinoderms: Filter feeders, like brittle stars, absorb nutrients in marine water. Suspension feeders use their arms to capture floating food particles. Grazers, like sea urchins, feed on both plants and animals, making them omnivores
Chaetognaths: predators of copepods, larval fish, crustaceans, and other chaetognaths
Hydrozoa: suspension feeders (plankton)
Tunicates: suspension feeders (plankton)
Sponges: filter feeders. detritus, plankton, viruses, bacteria, dissolved nutrients
Plankton: phytoplankton are primary producers and zooplankton eat phytoplankton
Mollusks: most graze algae or are filter feeders
Polychaetes: carnivorous, detritus, filter and deposit feeders

**Definition of diet classification categories.**
Primary consumer: feed on algae and plants.
Secondary consumer: eat herbivorous. In this case would be only insects and plankton.
Tertiary/Quaternary consumer: eat filter and suspension feeders (plankton), and omnivorous.
Parasites: species that feed on other but od not necessarily kill them.
